# A FRET based biosensor for measuring Gα13 activation in single cells

**DOI:** 10.1101/220632

**Authors:** Marieke Mastop, Nathalie R. Reinhard, Cristiane R. Zuconelli, Fenna Terwey, Theodorus W. J. Gadella, Jakobus van Unen, Merel J. W. Adjobo-Hermans, Joachim Goedhart

## Abstract

Förster Resonance Energy Transfer (FRET) provides a way to directly observe the activation of heterotrimeric G-proteins by G-protein coupled receptors (GPCRs). To this end, FRET based biosensors are made, employing heterotrimeric G-protein subunits tagged with fluorescent proteins. These FRET based biosensors complement existing, indirect, ways to observe GPCR activation. Here we report on the insertion of mTurquoise2 at several sites in the human Gα13 subunit. Three variants were found to be functional based on i) plasma membrane localization and ii) ability to recruit p115-RhoGEF upon activation of the LPA2 receptor. The tagged Gα13 subunits were used as FRET donor and combined with cp173Venus fused to the Gγ2 subunit as the acceptor. We constructed Gα13 biosensors by generating a single plasmid that produces Gα13-mTurquoise2, Gβ1 and cp173Venus-Gγ2. The Gα13 activation biosensors showed a rapid and robust response when used in primary human endothelial cells that were treated with thrombin, triggering endogenous protease activated receptors (PARs). This response was efficiently inhibited by the RGS domain of p115-RhoGEF and from the biosensor data we inferred that this is due to GAP activity. Finally, we demonstrated that the Gα13 sensor could be used to dissect heterotrimeric G-protein coupling efficiency in single living cells. We conclude that the Gα13 biosensor is a valuable tool for live-cell measurements that probe Gα13 activation.

## Introduction

G-protein coupled receptors (GPCRs) are members of a large family of membrane located receptors, with around 750 genes encoding a GPCR identified in the human genome (Vassilatis et al., 2003). These seven transmembrane containing proteins can perceive a wide variety of signals including light, hormones, ions and neurotransmitters (Wettschureck and Offermanns, 2005). GPCRs act as Guanine Exchange Factors (GEFs) (Rens-Domiano and Hamm, 1995) for heterotrimeric G-proteins. These protein complexes are comprised of a Gα, Gβ and Gγ subunit. The heterotrimer is a peripheral membrane protein complex due to lipid modification of the Gα and Gγ subunit (Wedegaertner et al., 1995).

The GEF activity is exerted on the Gα subunit, which can be converted from an inactive GDP-bound state to an active GTP-bound state (Hepler and Gilman, 1992). The activation of the complex results in a conformational change and in some cases the dissociation of the Gα subunit from the Gβγ dimer (Frank et al., 2005; Levitzki and Klein, 2002; Wettschureck and Offermanns, 2005). Both the activated GTP-bound Gα subunit and Gβγ dimer are capable of activating downstream effectors (Wettschureck and Offermanns, 2005).

Almost twenty different Gα subunits can be discerned and these are grouped in four classes; Gi/o, Gs, Gq and G12/13 (Wettschureck and Offermanns, 2005). Throughout this manuscript we will use Gα13 to indicate the subunit and G13 to indicate the heterotrimer, consisting of Gα13, Gβ and Gγ, the same terminology will be used for Gq/Gαq and Gi/Gαi, respectively. Each class of Gα subunits activates different downstream effectors (Hepler and Gilman, 1992). The best characterized effectors of the Gα12/Gα13 subunits are RhoGEFs that activate RhoA, e.g. LARG, PDZ-RhoGEF and p115-RhoGEF (Aittaleb et al., 2010; Siehler, 2009). For quite some time, it was thought that GPCR activation of the Gα12/ Gα13 subunits was the predominant way to activate RhoA. This view has changed over the last decade with the identification of RhoGEFs that can be activated by Gαq (Aittaleb et al., 2010; Rojas et al., 2007). Nowadays, it is clear that both Gαq and Gα12/ Gα13 can rapidly activate RhoA signaling in cells (Reinhard et al., 2017; van Unen et al., 2016c), albeit by activating different effectors. Since the Gq class and G12/13 class both efficiently activate RhoA, it has been difficult to distinguish which of these two heterotrimeric G-protein complexes is activated when only downstream effects are measured. To further complicate matters, GPCRs that activate G12/13 often activate Gq as well (Riobo and Manning, 2005; Worzfeld et al., 2008). Yet, it is clear that signaling through either Gq or G12/13 has different physiological effects (Takefuji et al., 2012; Wirth et al., 2008; Worzfeld et al., 2008). Therefore, it is necessary to have tools that can measure activation of the heterotrimeric G-protein itself.

Direct observation of heterotrimeric G-protein activation has classically been performed by quantifying the binding of radiolabeled nucleotides (Milligan, 2003). This approach is labor-intensive, uses disrupted cells and lacks temporal resolution. Moreover, this technique is generally less suitable for Gq, Gs and G12/13, due to their low expression levels compared to Gi (Strange, 2010). On the other hand, optical read-outs, often based on FRET and Bioluminescence Resonance Energy Transfer (BRET) techniques are well suited to measure signaling activity with high temporal resolution in intact cells (Lohse et al., 2012; van Unen et al., 2015). Several groups have generated BRET or FRET based biosensors for detecting events immediately downstream of activated GPCRs (Clister et al., 2015; Hébert et al., 2006; Lohse et al., 2012; Salahpour et al., 2012; van Unen et al., 2016b). Optical biosensors that are based on heterotrimeric G-proteins are particularly suited to report on GPCR activation (Janetopoulos and Devreotes, 2002). However, only a few optical biosensors for reporting activation of Gα12 or Gα13 have been reported. Sauliere et al. reported a BRET based biosensor for detection of activation of Gα13 activation, enabling the detection of biased-agonism via the angiotensin II type 1 receptor (AT1R) (Saulière et al., 2012). The approach, however, lacks spatial resolution. Improved luciferases, such as Nanoluciferase, enabled single cell BRET measurements, but longer acquisition times are required, so temporal resolution is decreased (Goyet et al., 2016). In general, FRET-based sensors have higher emission intensities, requiring shorter acquisition times to obtain sufficient spatial and temporal resolution. Thus, development of a FRET based biosensor reporting on the activation of Gα13 would be a real asset for GPCR signaling research. We have previously reported on single plasmid systems that enable the expression of a multimeric FRET based sensor for Gαq and Gαi (Goedhart et al., 2011; van Unen et al., 2016a). Here, we report on the development, characterization and application of a single plasmid, FRET based biosensor for the activation of Gα13.

## Results

### Strategy for tagging Gα13 with a fluorescent protein

To directly measure the activation of Gα13 with high spatiotemporal resolution in living cells, we aimed at generating a functional, fluorescent protein (FP) tagged Gα13 subunit. Gα subunits cannot be tagged at the N- or C-terminus since these are required for interaction with the Gβγ subunit and the GPCR (Wall et al., 1995). To functionally tag Gα13, the FP should be inserted in the Gα13 sequence, as was previously done for other Gα isoforms (Gibson and Gilman, 2006; Janetopoulos and Devreotes, 2002).

Initially, we used a sequence alignment of the four classes of Gα-proteins and identified the residues of Gαq (Adjobo-Hermans et al., 2011) and Gαi (van Unen et al., 2016a) after which we had previously inserted mTurquoise2 (figure 1B, supplemental figure S1). Based on sequence homology we chose to insert mTurquoise2 after residue Q144 of human Gα13, indicated with a black rectangular box in the alignment. Upon transfection of the plasmid encoding this variant, we observed cytoplasmic fluorescence. This localization probably reflects incorrect folding or targeting of the Gα subunit, since well-folded and functionally tagged variants are located at the plasma membrane (Adjobo-Hermans et al., 2011; Gibson and Gilman, 2006). Inspection of the crystal structure (PDB ID: 1ZCB), revealed that Q144 is part of an α- helix (αB2), which is likely to be disrupted after modification or insertion (Kreutz et al., 2006).

**Figure 1.**
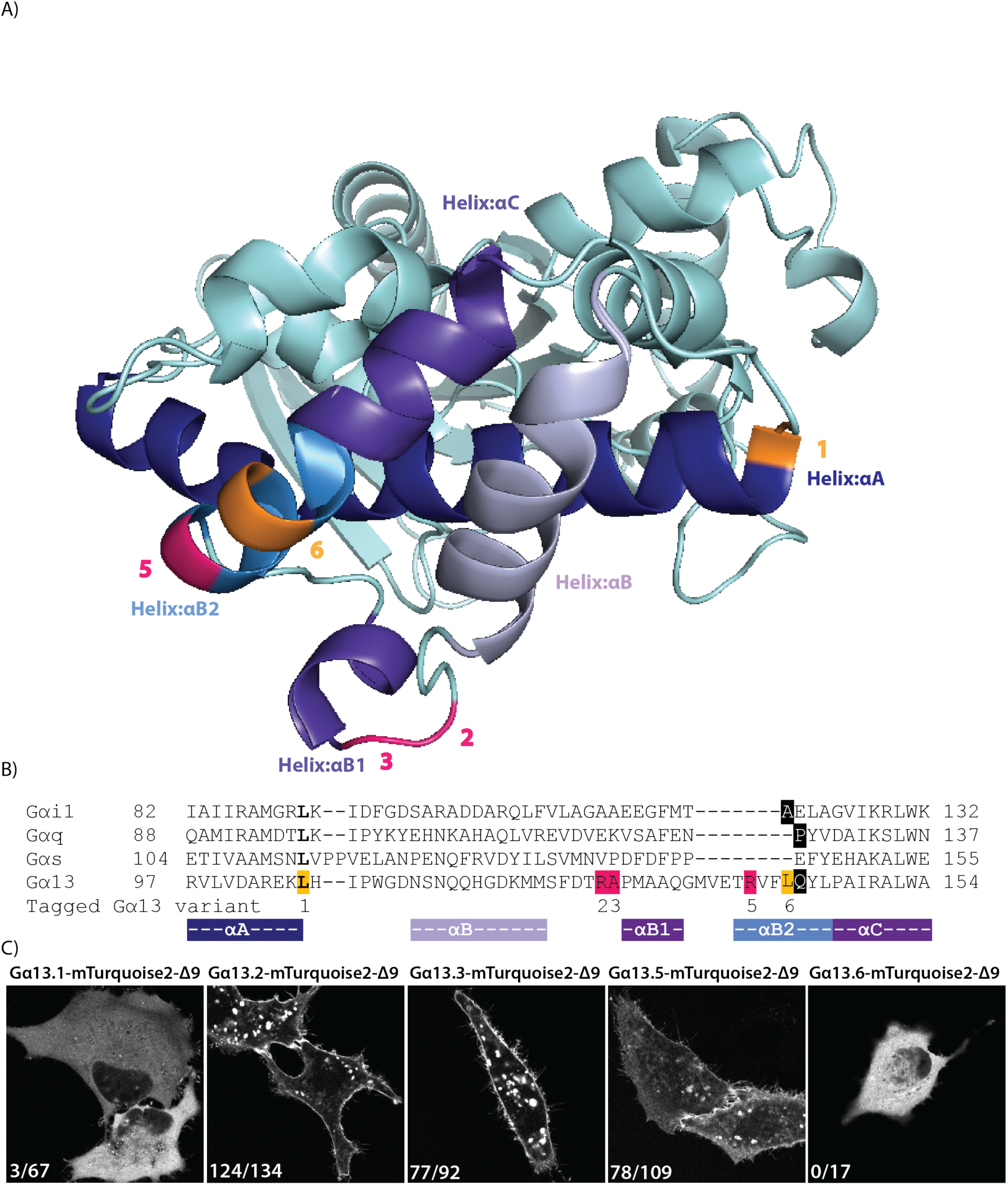
Insertion of a fluorescent protein at different positions in Gα13. (A) The protein structure of human Gα13 (PDB ID: 1ZCB). The highlighted residues indicate the amino acid preceding the inserted fluorescent protein. Successful sites for inserting mTurquoise2-Δ9 into Gα13 in pink and unsuccessful sites in orange. (B) A partial protein sequence alignment (full alignment see supplemental figure S1) of different Gα classes. The highlighted residues indicate the amino acid preceding the inserted fluorescent protein (or luciferase). In bold, the sites that were previously used to insert Rluc (Saulière et al., 2012). Insertion of mTurquoise2-Δ9 in Gα13 after residue Q144 (black) was based on homology with previous insertions in Gαq and Gαi (black). Successful sites for inserting mTurquoise2-Δ9 (R128, A129 and R140) in pink and unsuccessful sites (L106 and L143) in orange. The numbers indicated below the alignment correspond with the Gα13 variant numbers, used throughout the manuscript. The colors under the alignment match with the colors of the αHelices shown in (A). (C) Confocal images of the tagged Gα13 variants transiently expressed in HeLa cells. The numbers in the left bottom corner of each picture indicate the number of cells that showed plasma membrane localization out of the total number of cells analyzed. The width of the images is 76μm.

Next, we used the protein structure to select a number of residues that were nearby previous insertion sites and next to or close to the end of an α-helix (αA, αB2 or αB1). We also took along an insertion site (L106) that was previously used to insert a luciferase into Gα13 (Saulière et al., 2012). We used a truncated mTurquoise2, deleting the last 9 amino acids, since this worked well in the Gi sensor (van Unen et al., 2016a). The insertion sites are highlighted on the protein structure and in the sequence alignment in figure 1A,B. The different variants are numbered as Gα13.1, Gα13.2, Gα13.3, Gα13.5 and Gα13.6 throughout the manuscript.

The plasmids encoding the different tagged variants were transfected into HeLa cells and we observed striking differences in localization. As shown in figure 1C, variant 1 and 6 showed cytoplasmic localization. In contrast, strong plasma membrane labeling was observed for variant Gα13.2, Gα13.3 and Gα13.5. Since native Gα13 is expected to localize at the plasma membrane by virtue of palmitoylation (Wedegaertner et al., 1995), we decided to continue with the optimization and characterization of variants 2, 3 and 5.

### Functionality of the tagged Gα13 variants

The correct localization of the tagged Gα13 variants does not necessarily reflect functionality with respect to activity in signaling, i.e. the capacity to exchange GDP for GTP. To determine functionality, we turned to a dynamic cell-based assay. This assay is based on the observation that ectopic Gα13 expression is required for p115-RhoGEF relocation to the plasma membrane upon GPCR stimulation (Meyer et al., 2008). To evaluate the functionality of Gα13, we co-expressed the LPA2 receptor-P2A-mCherry (van Unen et al., 2016b), p115-RhoGEF, tagged with SYFP1, and different variants of Gα13, including an untagged, native variant (figure 2, supplemental figure S2). Cells in which Gα13 was not over-expressed did not show p115-RhoGEF relocation after GPCR activation (figure 2B,C). However, in the presence of native Gα13, a relocation of p115-RhoGEF was noticed (figure 2B,C). These findings are in agreement with previous findings and show that a functional Gα13 is required for the recruitment of p115-RhoGEF to the plasma membrane. Next, similar experiments were performed in the presence of Gα13 variants 2, 3 and 5, tagged with mTurquoise2Δ9. In all three cases a robust relocation of p115-RhoGEF was observed (figure 2). In contrast, the relocation was not evident when a Gα13 variant (variant 1) was employed that did not show efficient plasma membrane localization (figure 2B,C). Hence, we observed a correlation between membrane localization and functionality in the recruitment assay. Altogether, the results support the notion that the tagged variants 2, 3 and 5 of Gα13 can be activated by a GPCR and are capable of recruiting p115-RhoGEF.

**Figure 2.**
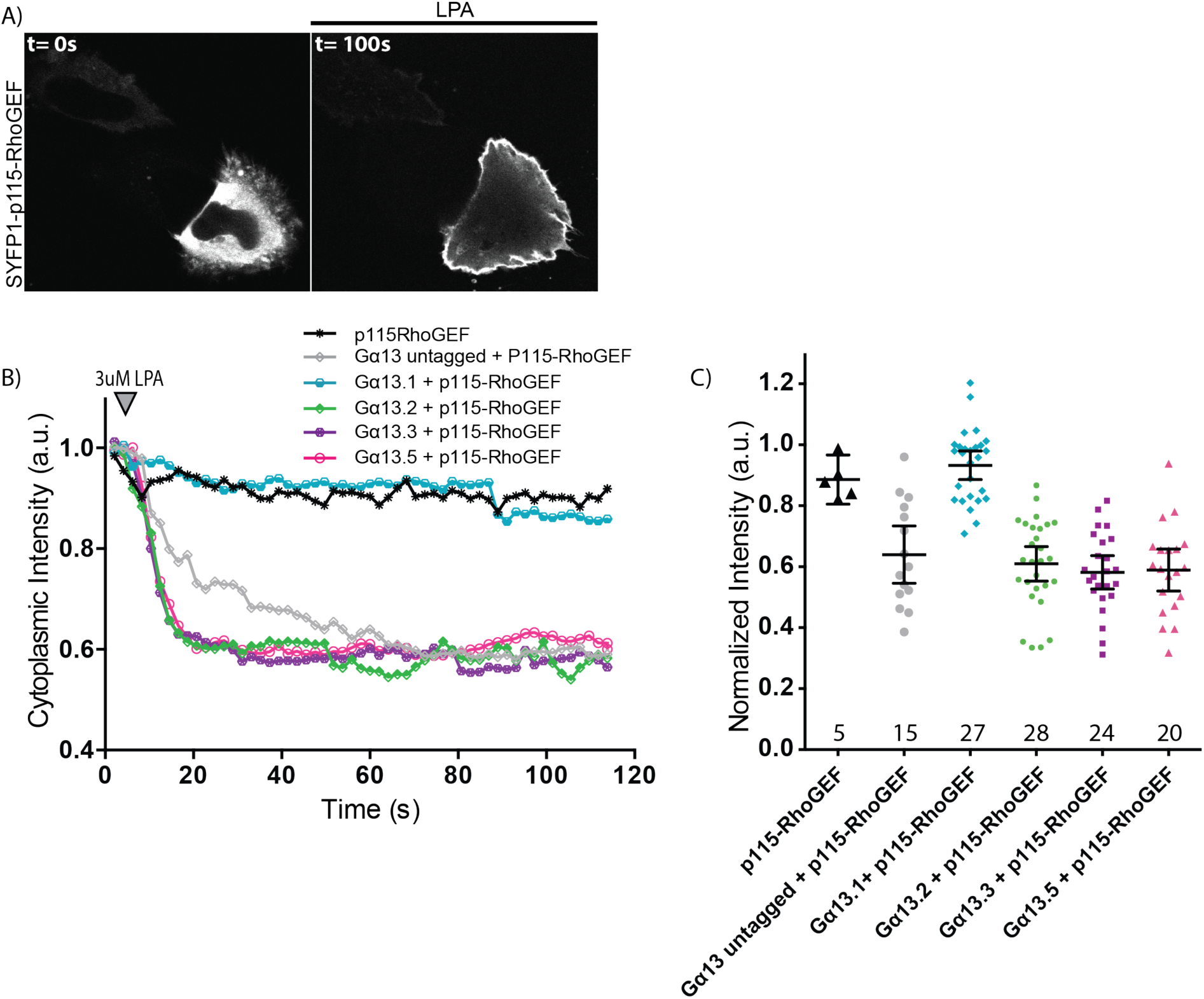
Capacity of the tagged Gα13 variants to recruit p115-RhoGEF. (A) Confocal images of a representative HeLa cell expressing SYFP1-p115-RhoGEF, Gα13.2-mTurquoise2-Δ9 and LPA2-P2A-mCherry (here only SYFP1-p115-RhoGEF is shown, for the localisation of the other constructs see supplemental figure S2) (before (t=0s) and after (t=100s) addition of 3µM LPA). The width of the pictures is 67μm. (B) The mean cytoplasmic fluorescence intensity of SYFP1-p115-RhoGEF over time. After 8s, 3μM LPA was added. All cells transiently expressed LPA2 receptor-P2A-mCherry. The number of cells imaged is p115-RhoGEF *n*=5, Gα13 untagged + p115-RhoGEF *n*=15, Gα13.1 + p115-RhoGEF *n*=27, Gα13.2 + p115-RhoGEF *n*=28, Gα13.3 + p115-RhoGEF *n*=24, Gα13.5 + p115-RhoGEF *n*=20. Data have been derived from three independent experiments. (C) Quantification of the fluorescence intensity at t==50s for each Gα13 variant, relative to t==0s. The dots indicate individual cells and the error bars show 95% confidence intervals. The numbers of cells analyzed is the same as in (B).

### Development and evaluation of Gα13 FRET based biosensors

Having engineered several correctly localizing Gα13 variants capable of recruiting p115-RhoGEF, we examined whether these variants can be used to report on G13 activation.

Using a FRET-based approach (as described for Gq (Adjobo-Hermans et al., 2011)), we monitored the interaction between the different generated mTurquoise2-tagged Gα13 constructs and a Gβγ dimer, consisting of untagged-Gβ and 173cpVenus- Gγ, on separate plasmids. These constructs allow acceptor (173cpVenus) – donor (mTurquoise2) FRET ratio measurements, where G13 activation results in a change in distance and/or orientation between the FRET pair, thereby inducing a FRET ratio decrease. Here we co-expressed an untagged LPA2-receptor, with one of the three tagged Gα13 variants, untagged Gβ, and tagged Gγ. Upon LPA2-receptor activation, a FRET ratio change was observed for all three selected Gα13 variants (figure 3A). The Gα13.2 variant showed the largest FRET ratio change and the Gα13.5 variant the lowest (figure 3A). Some cells expressing the Gα13.3 variant showed a slower response (figure 3A).

**Figure 3.**
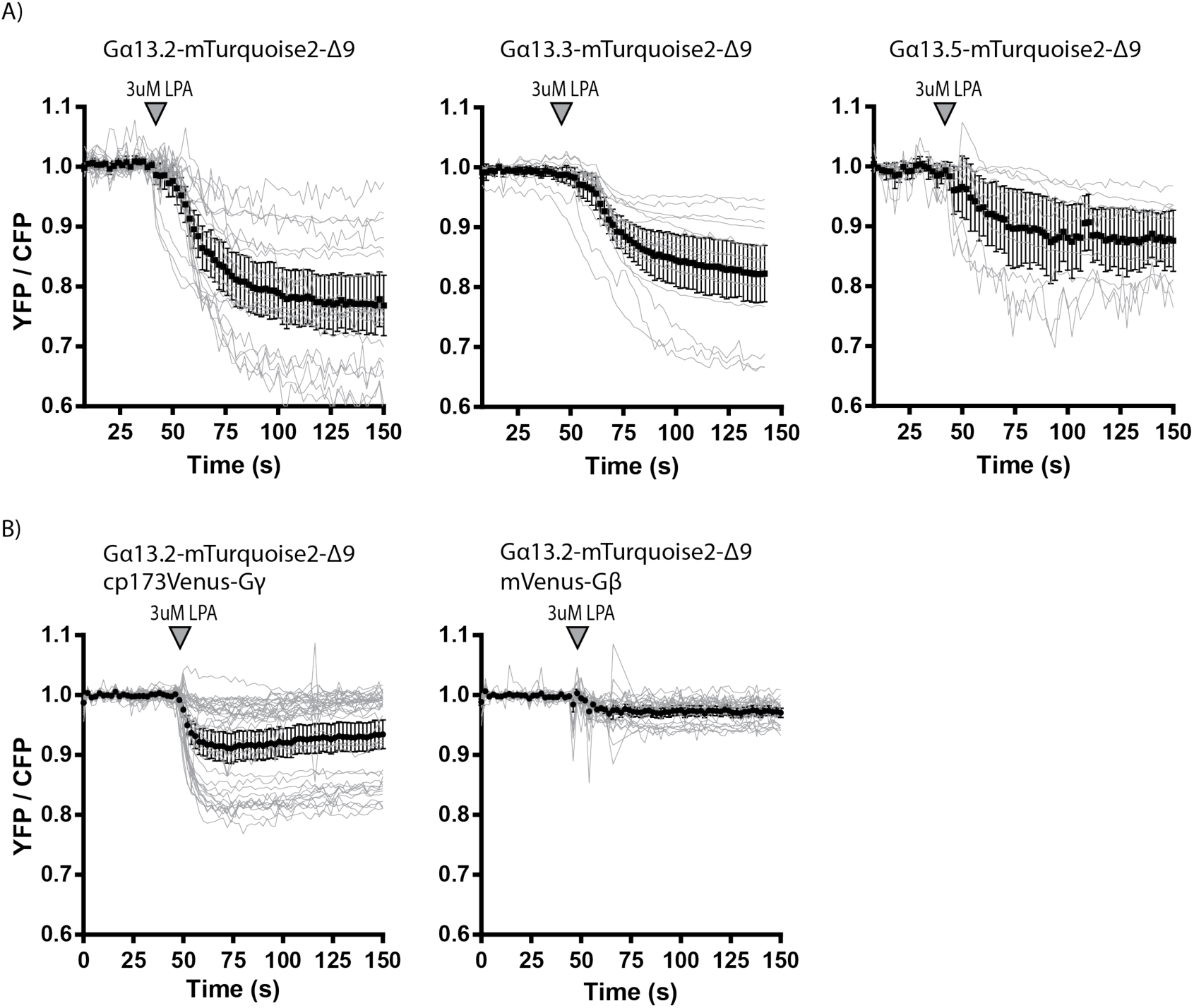
The ability of the tagged Gα13 variants to report on dynamic Gα13 activation via ratiometric FRET imaging. (A) Ratiometric FRET traces of HeLa cells expressing the LPA2 Receptor (untagged), Gβ (untagged), one of the Gα13 variants (as indicated in the title of the graphs) tagged with mTurquoise2-Δ9 and Gγ tagged with cp173Venus. The grey lines represent individual cells and the black graph represents the average of which the error bars indicate the 95% confidence intervals. LPA was added at t==42-50s, indicated by the arrowhead. The number of cells analyzed is: Gα13.2-mTurquoise2-Δ9 *n*=20, Gα13.3-mTurquoise2-Δ9 *n*=16 and Gα13.5-mTurquoise2-Δ9 *n*=11. (B) Ratiometric FRET traces of HeLa cells expressing the LPA2 Receptor (untagged), Gα13.2-mTurquoise2-Δ9 and either Gγ tagged with cp173Venus (*n*=38) (and untagged Gβ) or Gβ tagged with mVenus (*n*=25) (and untagged Gγ). LPA was added at t=50s, indicated by the arrowhead.

Recently, a FRET sensor for Gα13 was reported that employed a tagged Gα and a Gβ subunit (Bodmann et al., 2017). Therefore, we also examined if tagging the Gβ subunit instead of the Gγ subunit yields a higher FRET ratio change for the best performing Gα13 variant. From figure 3B can be inferred that tagging the Gγ subunit with the FRET acceptor results in the highest FRET ratio change upon activation of the G13 via the LPA2 receptor, which is consistent with our previous observations (Adjobo-Hermans et al., 2011). We have previously shown for Gαq (Goedhart et al., 2011) and Gαi (van Unen et al., 2016a) activation biosensors that a single expression plasmid ensures robust co-expression of the sensor components and simplifies the transfection. Since all three tagged Gα13 variants are able to report on Gα13 activation using 173cpVenus-Gγ as FRET acceptor, we developed single plasmid sensors using each of the Gα13 variants. Figure 4A shows a schematic overview of the plasmid design for a Gα13 sensor.

**Figure 4.**
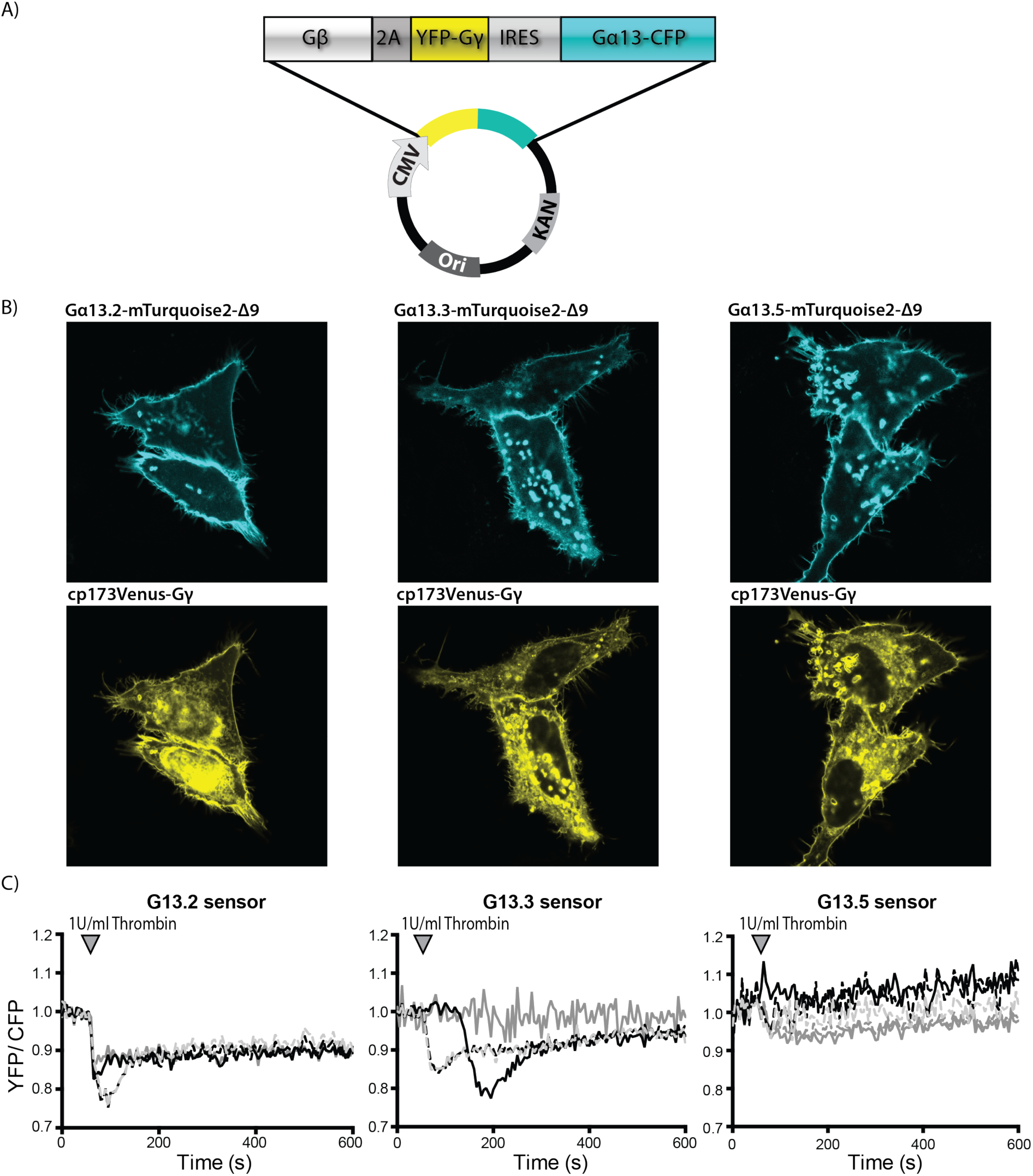
Development and characterization of Gα13 activation FRET based biosensors. (A) Architecture of the Gα13 biosensor construct, encoding Gβ-2A-cp173Venus-Gγ2-IRES-Gα13-mTurquoise2-Δ9, under control of the CMV promoter. (B) Confocal images showing the localization of the Gα13 in the sensor variants (upper, cyan) and cp173Venus-Gγ2 (lower, yellow) in HeLa cells. The width of the images is 75μm. (C) FRET ratio traces of HUVECs expressing the different G13 biosensors, stimulated with Thrombin at t==55s. Each line represents an individual cell. The number of cells analysed is: G13.2 sensor *n*=4, G13.3 sensor *n*=4, G13.5 sensor *n*=5.

To verify co-expression, the plasmids were introduced in HeLa cells. As can be inferred from figure 4B, the subcellular localization of the different Gα13 sensors expressed in HeLa cells is similar. The Gα13 variants are mainly located at the plasma membrane and Gγ is located to the plasma membrane and endomembranes as published (Goedhart et al., 2011).

Next, we evaluated the performance of the sensors in Human Umbilical Vein Endothelial cells (HUVEC). HUVECs are known to respond to thrombin, activating endogenous Protease Activated Receptors (PARs), which results in Gα13-RhoA signaling (Reinhard et al., 2016; Reinhard et al., 2017).

We observed no effect of ectopic sensor expression. When thrombin was added to HUVECs, the Gα13.2 sensor showed the most pronounced FRET change (Figure 4C). Based on the FRET ratio imaging data in HUVECs and HeLa, we selected the Gα13.2 sensor as the Gα13 activation biosensor of choice due to its high sensitivity and robust FRET ratio change upon Gα13 activation.

### Characterization of Gα13 inhibition by a GTPase Activating Protein

Our data show that the Gα13.2 sensor is sensitive enough to detect G13 signaling activated by endogenous thrombin receptors in HUVECs. We and others (Kelly et al., 2006; Martin et al., 2001; Reinhard et al., 2017) have used a Regulator of G-protein Signaling (RGS) domain of p115-RhoGEF as an inhibitor of G13 signaling. The RGS domain exhibits GTPase Activating Protein (GAP) activity (Kozasa et al., 1998).

However, it is unclear whether the inhibition by the RGS domain in cells is due to GAP activity or due to competitive binding of the RGS domain and downstream effectors to Gα13. In the latter case, we would expect a change in FRET ratio in the presence of the RGS domain. To gain insight in the mechanism of action, we employed the Gα13.2 sensor and co-expressed a membrane bound RGS (Lck-mCherry-p115-RGS), which is shown to effectively inhibit RhoA activation (Reinhard et al., 2017). In the presence of the RGS domain, we did not observe a FRET ratio change of the Gα13.2 sensor after adding thrombin to HUVECs. In the control sample (Lck-mCherry) we did observe a ratio change of the Gα13.2 sensor induced by thrombin (figure 5A,B), while resting state FRET ratios were similar for both conditions (data not shown). The lack of a FRET response in presence of the RGS domain, reflecting suppression of active GTP-bound Gα13, provides evidence that GAP activity is involved in the inhibitory effect of the RGS domain (figure 5A,B). Additionally, we looked into thrombin-induced contraction of HUVECs and show that over-expression of the RGS domain prevents cell contraction, even leading to an overall increase in cell area (figure 5C).

**Figure 5.**
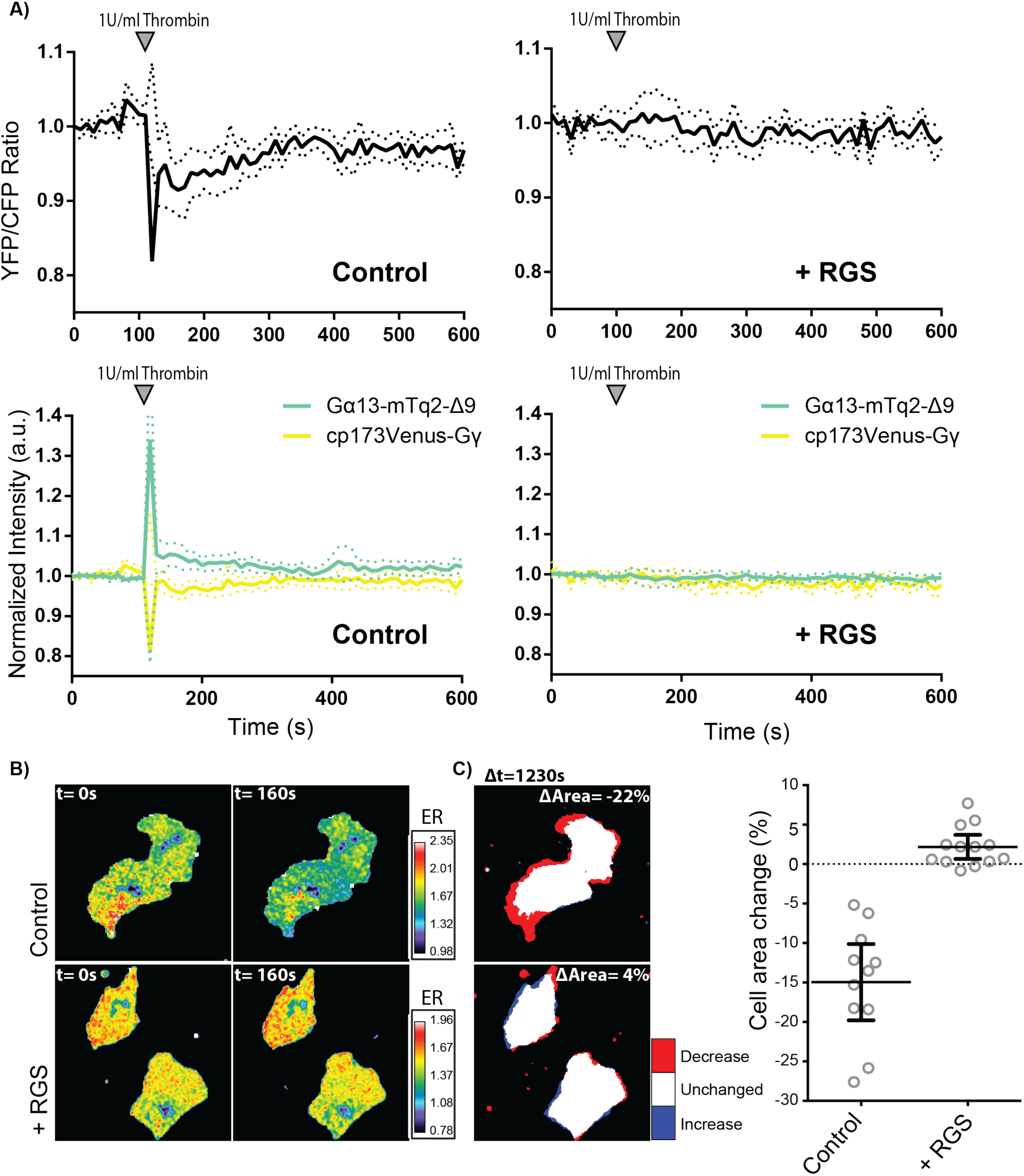
Effects of the p115-RhoGEF RGS domain on Gα13.2 activity and cell morphology. (A) Normalized ratiometric traces (upper graphs) and corresponding YFP and CFP traces (lower graphs) (dotted lines depict 95% CI) of HUVECs that were transfected with either the Gα13.2 FRET sensor and Lck-mCherry (Control, *n*=11) or the Gα13.2 FRET sensor and Lck-mCherry-RGS (+ RGS, *n*=13). Cells were stimulated at t = 110s. (B) Ratiometric images of representative cells measured in (A). Cool colors represent low YFP/CFP ratios, corresponding to emission ratios (ERs) on the right. (C) Cell area change of the cells measured in (B), visualized according to the LUT panel on the right. Dotplots on the right represent individual measurements (± 95% CI) of corresponding cells measured in (A). Image width = 54μm.

### Application of the Gα13.2 biosensor in GPCR activation assays

Published data on GPCRs coupling to Gα13 is often based on indirect measures and since downstream signaling effects overlap with the downstream signaling effects of Gq, it is difficult to draw solid conclusions about the involvement of either subunit (Riobo and Manning, 2005). Our novel FRET sensor enables direct observation of Gα13 activation. Therefore, we evaluated the Gαq and Gα13 responses to the stimulation of a selection of GPCRs published as coupling to Gα13 (Inoue et al., 2012; Meyer et al., 2008; Navenot et al., 2009; Saulière et al., 2012).

These experiments were performed in HeLa cells transiently expressing the corresponding GPCR. As a control, cells only expressing the Gα13 or Gαq sensor were stimulated, which did not elicit a noticeable sensor response (figure 6A). Upon LPA2 receptor (LPA2R) stimulation, Gαq and Gα13 were both activated, which corresponds to published data. Of note is that a number of cells (47 out of 60 for Gαq and 23 out of 37 for Gα13), transiently expressing the LPA2 receptor, failed to show a Gα13 or Gαq response (figure 6B). Upon Angiotensin II type 1 receptor (AT1R) stimulation an obvious response of the Gαq sensor is observed compared to a small, but consistent response of the Gα13 sensor (figure 6C). The Kisspeptin receptor (KissR or GPR54) showed a clear Gαq and no Gα13 sensor response upon stimulation (figure 6D). Together, these results indicate that LPA2R and to a lesser extend AT1R are coupled to Gα13, while the KissR is only coupled to Gαq and not to Gα13. Moreover, it shows that our novel Gα13 sensor can indeed be used to distinguish between Gα13- and Gαq-coupled GPCR signaling.

**Figure 6.**
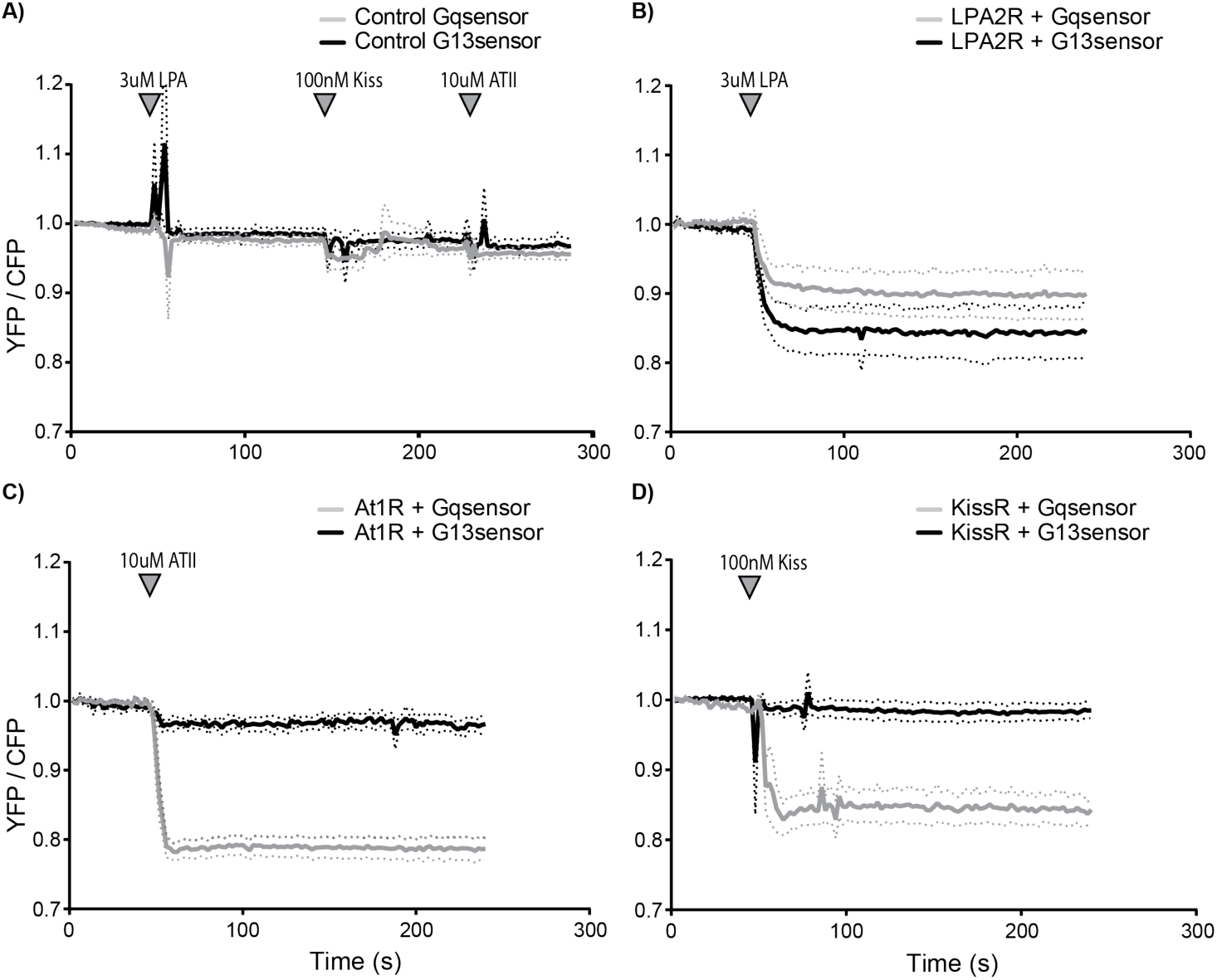
Direct observation of Gα13 and Gαq activation by different GPCRs. Normalized ratio-metric FRET traces of HeLa cells transfected with the Gq sensor (grey line) or the G13.2 sensor (black line) (dotted lines depict 95% CI). (A) As a control, cells expressing only the Gq (*n*=37) or G13.2 (*n*=20) sensor were measured. Agonists were sequentially added after 50s, 150s and 230s of imaging. (B) Ratio traces of cells transfected with an untagged LPA2 receptor next to the Gaq (*n*=13 (out of 60 in total)) or the Gα13.2 (*n*=14 (out of 37 in total)), stimulated at t=50s. (C) Ratio traces of cells transfected with angiotensinII type 1 receptor-P2A-mCherry next to the Gaq (*n*=22) or the Gα13.2 (*n*=9) sensor, stimulated at t=50s. (D) Ratio traces of cells transfected with an untagged kiss-receptor next to the Gαq (*n*=13) or the Gα13.2 (*n*=30) sensor, stimulated at t=50s (indicated with the arrowhead).

## Discussion

The combined activation of different heterotrimeric G-protein classes by a GPCR defines which signaling networks will be activated. For one of the classes of G-proteins, G12/13, it has been notoriously difficult to measure its activation. Here, we report on the development, characterization and application of a FRET based biosensor for Gα13 activation. This novel tool can be used to study the activation of Gα13 by GPCRs in single living cells. We furthermore show that the G13 sensor is sensitive enough to report on the activation of an endogenous receptor in human primary cells (HUVECs).

Thus far, the tools available to study the activation of Gα13 in single living cells have been limited. BRET-based strategies have been used to study activation of the G12 class. To this end, *Renilla reniformis* luciferase (Rluc) was inserted in Gα13 after residue Ile 108 (Ayoub et al., 2010) or Leu 106 (Saulière et al., 2012). However, we find that insertion of mTurquoise2 after residue Leu 106 (Gα13.1) did not result in a functionally tagged subunit, possibly because Leu106 is part of an α-helix.

Three of the five evaluated mTurquoise2 insertion sites resulted in plasma membrane localized, tagged Gα13, which were able to recruit p115-RhoGEF to the plasma membrane. The cytoplasmic-localized, tagged Gα13.1 variant was not able to recruit p115-RhoGEF, indicating that plasma membrane localization is required for functionality. Since all functional Gα13 variants could report on G13 activation as determined by FRET ratio-imaging, we developed single plasmid sensors for these three variants. In HeLa cells the localization is as expected, Gα13 at the plasma membrane and Gγ at the plasma membrane and endomembranes. The Gα13 activation biosensors were expressed in HUVECs and could report on G13 activation via endogenous Protease Activated Receptors (PARs). The Gα13.2 sensor is the best performing Gα13 biosensor, based on its sensitivity and magnitude of the FRET ratio change upon activation. Further improvement of the sensors might be achieved by varying FRET pairs (Mastop et al., 2017) or changing their relative orientation by circular permutation (Fritz et al., 2013).

While our manuscript was in preparation, another mTurquoise2 tagged Gα13 variant was reported (Bodmann et al., 2017). In this study, the fluorescent protein was inserted after residue 127 of Gα13, closely resembling our Gα13.2 variant, and it showed plasma membrane localization. Moreover, Bodmann et al. observed a FRET change when an YFP tagged Gα13 variant was used in combination with a CFP tagged Gβ subunit to report on the activation of the thromboxane A2 receptor (Bodmann et al., 2017). The independent observation of a FRET change using a similar tagged Gα13 variant, supports our notion that the G13.2 FRET based biosensor is a valuable tool for studying the activation of Gα13.

An advantage of our FRET sensor is that a higher ratio change is detected than for the sensor reported by Bodmann et al. 2017. While they detected a maximal FRET ratio change of 10%, we could reach a FRET ratio change of up to 30% (figure 3). This might be due to using a tagged Gβ instead of Gγ subunit, since we also observe a lower FRET ratio change when a tagged Gβ is employed (figure 3B). It is remarkable that Bodmann et al. detect a FRET increase upon Gα13 stimulation (increased YFP, decreased CFP intensity) whereas we detect a FRET decrease upon Gα13 stimulation (figure 3B). All of the G-protein sensors reported by us (Gαq, Giα1,2,3 and Gα13) show a FRET decrease upon activation under all conditions tested thus far (Adjobo-Hermans et al., 2011; Reinhard et al., 2017; van Unen et al., 2016a), consistent with a dissociation of Gα and Gβγ subunits. Moreover, Gα13 subunits, tagged at different sites (Gα13.2, Gα13.3 and G*α*13.5) all show a FRET decrease (figure 3). We do not have an explanation for the opposite changes in FRET efficiency observed by Bodmann et al. and by us (Bodmann et al., 2017).

Another advantage of the sensor that we constructed is that it enables the simultaneous production of the three proteins that comprise the heterotrimer from a single plasmid and that the donor and acceptor are present at a well-defined stoichiometry (Goedhart et al., 2011). This simplifies the distribution and application of the sensor. Especially in primary cells, such as HUVECs, it is more complicated to obtain simultaneous protein expression from multiple plasmids, whereas transient expression of the single sensor plasmid in HUVECs was efficient.

The RGS domain of p115-RhoGEF effectively inhibits GPCR-Gα12/13 mediated RhoA activation in endothelial cells (Reinhard et al., 2017) and cell contraction as shown in figure 5c. Under these conditions, the Gα13.2 sensor did not display a FRET ratio change, reflecting a lack of activation. The absence of a FRET change is consistent with the idea that the GAP activity of the RGS domain shuts down Gα13 activity (Kozasa et al., 1998).

Applying the Gα13.2 sensor in HeLa cells, we showed that Gα13 is activated via the LPA2 receptor and to a much lesser extend via the AT1 receptor, while the Kiss receptor only elicited a Gq sensor response. The GPCRs analyzed in this study, were reported as coupling to Gα13 and / or Gαq in other studies. Of note, transient expression of the LPA2 receptor shows several non-responders in terms of Gαq or Gα13 activity. This could be explained by insufficient levels of receptor to achieve detectable activation.

Strikingly, the Kiss receptor did not activate the Gα13 biosensor, indicating that aspecific, promiscuous coupling of a GPCR to an ectopically expressed G-protein biosensor is not detected. Conversely, we previously observed that HUVEC treated with S1P do show G13 activation, but no Gq activation (Reinhard et al., 2017). Together, these observations suggest that our FRET based biosensor toolkit provides a way to determine GPCR coupling selectivity towards heterotrimeric G-proteins in living cells. It would be advisable to complement such studies with more downstream read-outs to verify the possible perturbation by the ectopically expressed G-protein sensors.

In summary, the results obtained in this study point out that the Gα13.2 biosensor is a sensitive Gα13 activation sensor for live cell imaging, that is suitable for application in primary cells and able to detect endogenous GPCR activation.

## Material & Methods

### Cloning/plasmid construction

mTurquoise2-Δ9 was inserted in the Gα13 sequence using a previously reported strategy (van Unen et al., 2016a). A PCR was performed on the mTurquoise2 sequence to truncate it and flank it with AgeI sites using primers Fw 5’-ATACCGGTTCTATGGTGAGCAAGGGCG-3’ and Rv 5’-TAACCGGTGATCCCGGCGGC -3’. To determine at which positions we wanted to insert mTurquoise2 in Gα13, we used Pymol to look at the structure of Gα13 and we used ClustalW to make an alignment of multiple Gα classes. A whole-vector PCR was performed on a pcDNA vector encoding human Gα13 (ordered from cDNA.org) to introduce an AgeI restriction site at the spot where we wanted to insert mTurquoise2, using primers Fw 5’- ATACCGGTCATATTCCCTGGGGAGACAAC-3’ and Rv 5’- ATACCGGTAAGCTTCTCTCGAGCATCAAC-3’for Gα13.1, primers Fw 5’-ATACCGGTGCCCCCATGGCAGCCC-3’ and Rv 5’- ATACCGGTCCGGGTATCAAACGACATCATCTTATC-3’ for Gα13.2, primers Fw 5’- ATACCGGTCCCATGGCAGCCCAAGG-3’ and Rv 5’-ATACCGGTGGCCCGGGTATCAAACGAC-3’ for Gα13.3, primers Fw 5’-ATACCGGTGTTTTCTTACAATATCTTCCTGCTATAAGA-3’ and Rv 5’- ATACCGGTCCTTGTTTCCACCATTCCTTG-3’ for Gα13.5 and primers Fw 5’- ATCCGGTCAATATCTTCCTGCTATAAGAGCA-3’ and Rv 5’-ATACCGGTTAAGAAAACCCTTGTTTCCACC-3’ for Gα13.6. The Gα13 pcDNA vector including AgeI site and the mTq2 PCR product were cut with AgeI and ligated, resulting in mTurquoise2 tagged Gα13. Of note, the cloning of variant Gα13.4, with an insertion of mTurquoise2-Δ9 after T139, failed.

We used GFP-p115-RhoGEF (a kind gift of Keith Burridge, UNC, Chapel Hill, USA) as a template to amplify p115-RhoGEF with the primers Fw 5’- AACAGATCTCTTGGTACCGAGCTCGGATC-3’ and Rv 5’- AGCGTCGACTCAAGTGCAGCCAGGCTG-3’. The PCR product, flanked with the restriction sites BglII and SalI, was used to clone p115-RhoGEF into a clontech-style C1 vector, generating mVenus-p115-RhoGEF.

The untagged LPA2 receptor was ordered from cDNA.org. To create the clontech-style N1 LPA2receptor-P2A-mCherry construct, a PCR was performed using primers Fw 5’- AGGTCTATATAAGCAGAGC-3’ and Rv 5’- TATGTCGACTTGGGTGGAGTCATCAGTG-3’. The N1-P2A-mCherry construct, described previously (van Unen et al., 2016b) and the LPA2 receptor PCR product were digested with EcoRI and SalI and ligation resulted in the LPA2receptor-P2A-mCherry construct.

The G13 single plasmid sensor variants were constructed as descripted previously by overlap-extension PCR (Heckman and Pease, 2007; van Unen et al., 2016a). The first PCRs were performed on the previously published Gαq sensor (Goedhart et al., 2011) using primerA Fw 5’-GAAGTTTTTCTGTGCCATCC-3’ and primerB Rv 5’-GTCCGCCATATTATCATCGTGTTTTTCAAAG-3’ and on the mTurquoise2 tagged Gα13 variants using primerC Fw 5’-ACGATGATAATATGGCGGACTTCCTGC-3’ and primerD Rv 5’- ATCAGCGGGTTTAAACG-3’. The second PCR was performed on a mix of both PCR products using primerA and primerD. This second PCR product and the Gαq sensor were digested with SacI and XbaI and the PCR product was ligated into the sensor, resulting in a G13 single plasmid sensor.

LCK-mCherry-p115-RhoGEF-RGS was constructed as described before (Reinhard et al., 2017). The p115- RhoGEF-RGS domain (amino acid 1-252) was PCR amplified using Fw 5’- GAGATCAGATCTATGGAAGACTTCGCCCGAG-3’ and Rv 5’-GAGATCGAATTCTTAGTTCCCCATCACCTTTTTC-3’. The PCR product and clontech-style C1 LCK-mCherry vector were digested with BglII and EcoRI and ligation resulted in a clontech-style C1 vector encoding LCK-mCherry-p115-RhoGEF-RGS.

The untagged Kiss receptor was purchased from www.cDNA.org.

The rAT1aR-mVenus was a kind gift from Peter Várnai (Semmelweis University, Hungary). The coding sequence of AT1R was inserted into mCherry-N1 with a P2A peptide to obtain AT1R-P2A-mCherry.

### Cell culture and sample preparation

HeLa cells (CCL-2, American Tissue Culture Collection; Manassas,VA, USA) were cultured in Dulbecco’s modified Eagle’s medium (DMEM) (Gibco, cat# 61965–059) supplemented with 10% fetal bovine serum (Invitrogen, cat# 10270–106), 100U/ml penicillin and 100 μg/ml streptomycin at 37 °C in 7% CO_2_. For microscopy experiments, cells were grown on 24mm Ø round coverslips, 0.13–0.16 mm thick (Menzel, cat# 360208) to 50% confluency and transfected with 500ng plasmid DNA, 1 μL Lipofectamin 2000 (Invitrogen, cat# 11668–019) or 4.5 μl PEI (1 mg/ml) in water (pH 7.3) and 100 μl OptiMEM (Gibco, cat# 31985–047) per 35mm Ø dish holding a 24mm Ø coverslip. One day after transfection the coverslip was mounted in a cell chamber (Attofluor, Invitrogen). Microscopy medium (20 mM HEPES (pH = 7.4), 137 mM NaCL, 5.4 mM KCl, 1.8 mM CaCl2, 0.8 mM MgCl2 and 20 mM glucose) was added to the coverslip in the cell chamber. The ratiometric FRET experiments were performed at 37 °C.

Primary HUVECs, acquired from Lonza (Verviers, Belgium) were seeded on fibronectin (FN)-coated culture flasks. HUVECs, grown in EGM-2 medium, supplemented with singlequots (Lonza) at 37 °C in 5% CO_2_. For microscopy experiments HUVECs were transfected at passage #4 or #5 with a Neon transfection system (MPK5000, Invitrogen) and Neon transfection kit (Invitrogen) and grown on FN-coated 24mm Ø round coverslips, 0.13–0.16 mm thick (Menzel, cat# 360208). Per transfection, 2µg plasmid DNA was used and a single pulse was generated at 1300 Volt for 30 ms. (Reinhard et al., 2017). The ratiometric FRET experiments were performed at 37 °C in EGM-2 medium and in 5% CO_2_.

### Confocal microscopy

To obtain confocal images of live HeLa cells transiently expressing either a tagged Gα13 variant or G13 single plasmid sensor, a Nikon A1 confocal microscope, equipped with a 60x oil immersion objective (Plan Apochromat VC, NA 1.4), was used. The pinhole size was set to 1 Airy unit. To check the localization of tagged Gα13 variants, samples were excited with a 457nm laser line, a 457/514 dichroic mirror was used and the emission was filtered through a 482/35BP filter. To check the localisation of the G13 single plasmid sensor constructs, samples were excited with a 440nm (CFP) and a 514nm (YFP) laser line, a 457/514 dichroic mirror was used and the emission was filtered through a 482/35BP (CFP) or 540/30BP (YFP), respectively. Images were acquired with sequential line scanning modus, to avoid bleedthrough.

### p115-RhoGEF recruitment assay

For the p115-RhoGEF recruitment assay, HeLa cells were cultivated in RPMI medium supplemented with 10% fetal calf serum and L-Glutamine (2 mM) (PAN Biotech GmbH, Aidenbach, Germany) and kept at 37°C in a 5% CO_2_ atmosphere. Cells were harvested, and 50.000 cells/well were seeded in eight-well µ-slides (Ibidi). After 24 hours, cells were transiently transfected with 0.25 μg of the LPA2receptor-P2A-mCherry, the SYFP1-p115-RhoGEF and the Gα13 variants tagged with mTurquoise2 (Gα13.1, Gα13.2, Gα13.3, Gα13.5). Transfection of HeLa cells was performed using Lipofectamine 3000 and Plus Reagent, according to the manufacturer’s instructions (Invitrogen). Twenty-four hours after transfection, the growth medium was replaced by RPMI phenol red-free medium and measurements were performed after a total incubation time of 48h. Throughout the measurements, cells were kept at 37°C. Cells were stimulated with Oleoyl-L-α-lysophosphatidic acid (10 µM, Sigma) at the indicated time point. Confocal images were taken with a Leica TCS SP5 laser scanning confocal microscope (Leica Microsystems, Mannheim, Germany) equipped with an HCX PL APO 63×, N.A. 1.2, water immersion lens. mTurquoise2 was excited at 458 nm and emission was detected between 465-500 nm; SYFP1 was excited at 514 nm and emission was detected between 520-550 nm; mCherry was excited at 561 nm and emission was detected between 600-670 nm. To avoid bleedthrough, images were acquired in the sequential line scanning modus. Image analysis was performed with Fiji.

### Widefield microscopy

#### Ratiometric FRET imaging HeLa cells

Ratiometric FRET experiments were performed on a wide-field fluorescence microscope (Axiovert 200 M; Carl Zeiss GmbH)(Adjobo-Hermans et al., 2011) equipped with a xenon arc lamp with monochromator (Cairn Research, Faversham, Kent, UK) and Metamorph 6.1 software, for 240s or 288s (controls in figure 6A) and with a time interval of 2s. The fluorescence intensity of the donor and acceptor were recorded with an exposure time of 200ms per image using a 40x objective (oil-immersion Plan-Neo- fluor 40×/1.30; Carl Zeiss GmbH). HeLa cells were used, expressing cp173Venus-Gγ (or untagged Gγ in figure 3B), untagged Gβ (or mVenus tagged Gβ in figure 3B) and one of the mTurquoise2 tagged Gα13 variants from multiple plasmids or expressing the single plasmid Gq-sensor or one of the G13 sensor variants. A GPCR is expressed from a separate plasmid, which was either untagged LPA2 receptor, untagged Kiss receptor or angiotensinII type 1 receptor-P2A-mCherry. Fluorophores were excited with 420 nm light (slit width 30 nm), mTq2 emission was detected with the BP470/30 filter, YFP emission was detected with the BP535/30 filter and RFP emission was detected with a BP620/60 filter by turning the filter wheel. After 50 s HeLa cells (unless stated otherwise) were stimulated with either a final concentration of 3μM LPA (Sigma), 100nM Kiss-1 (112-121) Amide (Phoenix pharmaceuticals) or 10μM angiotensin (Sigma). The curves were normalized to the average intensity of the first 5 frames that were recorded. ImageJ was used to perform a background correction and calculation of mean intensity of each cell for each time point. Cells that did not show a visible response were not used for the analysis. The total number of cells imaged and the number of cells analyzed (“the responders”) are indicated in the figure legends.

#### Ratiometric FRET imaging HUVECs

To perform ratiometric FRET experiments on HUVECs we used the same microscopy equipment and filter settings as were used to perform ratiometric FRET imaging of HeLa cells, however there are some differences in the way the data is recorded. The HUVECs are imaged for 1230s with a time interval of 10s and the imaging is performed at 37 °C in EGM-2 medium and in 5% CO_2_. After 110s, HUVECs were stimulated with a final concentration of 1U/ml thrombin (Haematologic Technologies). The image processing procedure that was used to display the change in cell area during live cell microscopy of HUVECs is described elsewhere (Reinhard et al., 2017)). In order to show the FRET ratio in images of cells, imageJ was used. We first converted the CFP and the YFP stack to 32-bit type and corrected for background signal. Then the stacks were divided to obtain a YFP/CFP stack. We used the CFP stack to make a binary mask and multiplied this mask with the YFP/CFP stack. Finally, we applied a smooth filter to reduce noise and used the lookup table (LUT) to visualize changes in FRET ratio.

## Acknowledgments

We thank the members of our labs for their continuous interest and support of this project. The plasmid with rAT1aR-mVenus was a kind gift from Peter Várnai (Semmelweis University, Hungary) and GFP-p115-RhoGEF was provided by Keith Burridge (UNC, Chapel Hill, USA).

## Author contributions

M.M. N.R. C.R.Z. performed experiments and analyzed data

F.T. and J.v.U. were involved in designing and constructing plasmids.

M.J.W.A-H and J.G. contributed to the data analysis and interpretation

M.M. and J.G. wrote the manuscript.

T.W.J.G. assisted with experimental design and interpretation of data.

All authors approved the final manuscript.

## Data availability

Plasmids and plasmid information is available from addgene (http://www.addgene.org/Dorus_Gadella/), most experimental data is presented in the manuscript, and the remainder is available from the corresponding author upon request.

## Supplementary figures

**Supplemental figure S1.**
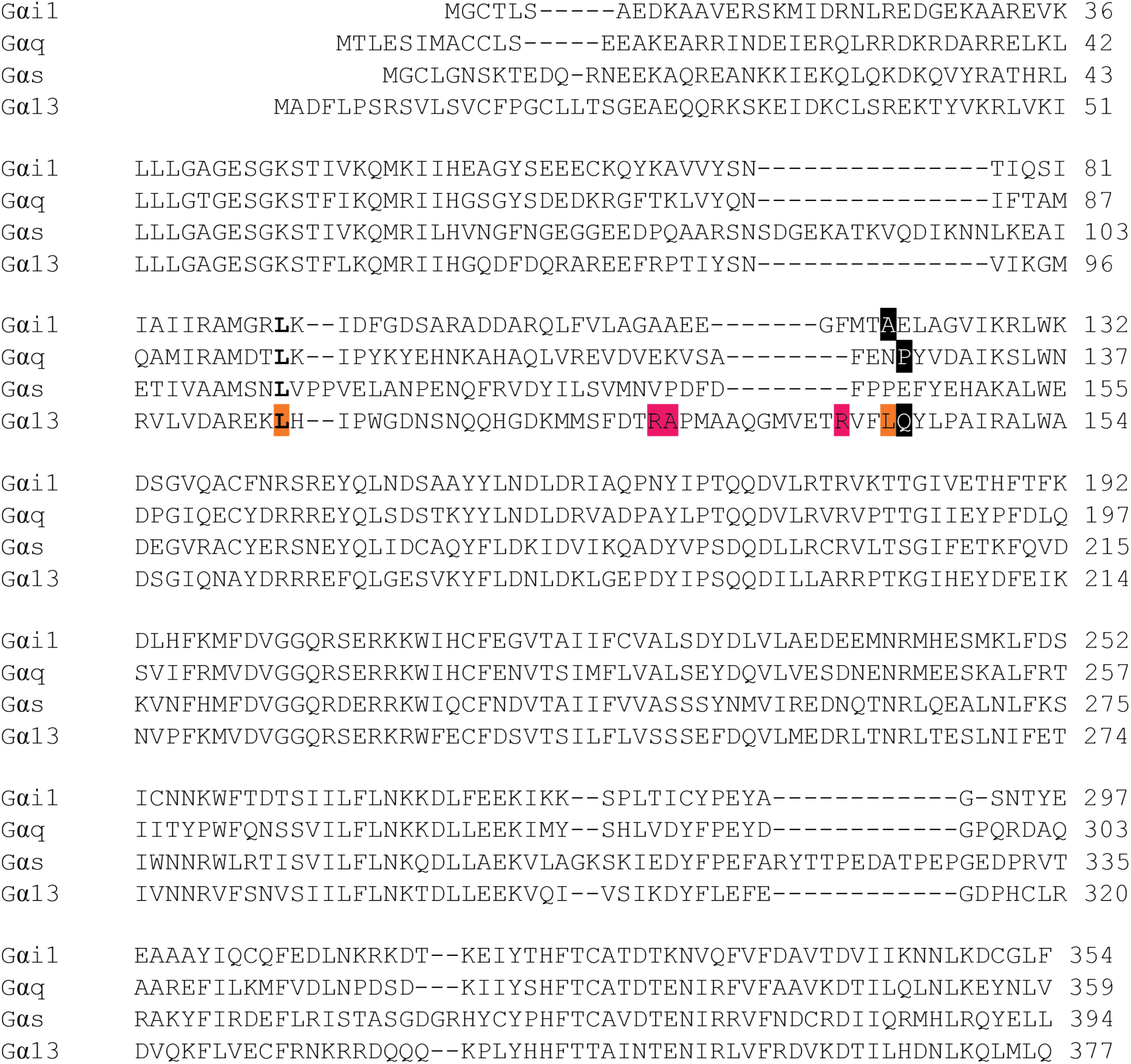
Full amino acid alignment of four classes of human Gα subunits of heterotrimeric G-proteins. Note that the amino acid sequence of Gαs is that of the short isoform. The highlighted residues indicate the amino acid preceding the inserted fluorescent protein (or luciferase). In bold, the sites that were previously used to insert Rluc (Saulière et al., 2012). Insertion of mTurquoise2-Δ9 in Gα13 after residue Q144 (black) was based on homology with previous insertions in Gαq and Gαi (black). Successful sites for inserting mTurquoise2-Δ9 (R128, A129 and R140) in pink and unsuccessful sites (L106 and L143) in orange.

**Supplemental figure S2.**
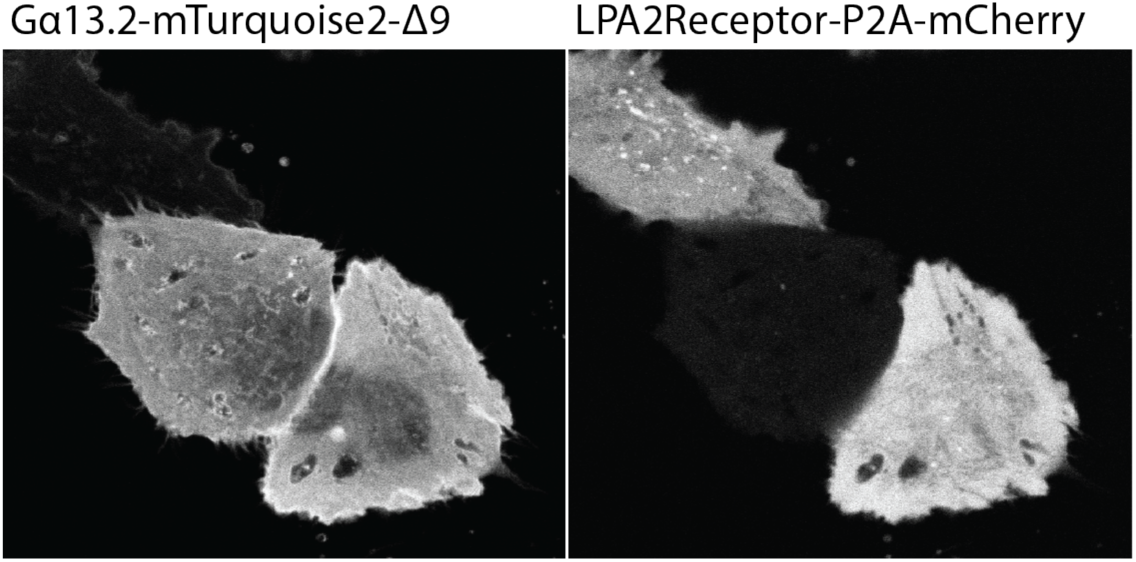
Confocal images of HeLa cells expressing both the Gα13.2- mTurquoise2 (left) and LPA2-p2A-mCherry (right) used in p115-RhoGEF recruitment assay (figure 2). The width of the images is 67μm.

